# ITSxRust: ITS region extraction with partial-chain recovery and structured diagnostics for long-read amplicon sequencing

**DOI:** 10.64898/2026.02.25.707950

**Authors:** Aaron O’Brien, Catalina Lagos, Kiara Fernández, Bárbara Ojeda, Pilar Parada

## Abstract

As long-read amplicon sequencing (e.g., Oxford Nanopore and PacBio HiFi) becomes routine for fungal metabarcoding, identifying and extracting ITS subregions at scale has become a throughput and robustness bottleneck. The nuclear ribosomal internal transcribed spacer (ITS) region is the formal DNA barcode for fungi and is widely used for taxonomic profiling of fungal communities [Schoch et al., 2012]. Standard preprocessing locates conserved ribosomal flanks with hidden Markov profile models (profile-HMMs) to extract ITS1, 5.8S, ITS2, or the full ITS, as implemented in ITSx [Bengtsson-Palme et al., 2013] and ITSxpress [Rivers et al., 2018, Einarsson and Rivers, 2024].

Here we describe ITSxRust, a Rust-based ITS extractor designed for long-read scale. ITSxRust coordinates HMMER searches with efficient Rust-native I/O and sequence processing, optionally reduces redundant searches via dereplication, provides ONT and HiFi parameter presets, and emits structured failure diagnostics and QC summaries. On an Oxford Nanopore ITS dataset (54,659 reads), ITSxRust extracted the full ITS region from 75.3% of reads, exceeding both ITSx (69.9%) and ITSxpress v2 (41.4%), while running 4.6× faster than ITSx. In addition, a partial-chain fallback strategy that extracts subregions using two-anchor pairs when the full four-anchor chain is unavailable recovered an additional 10,725 reads that would otherwise be discarded.

## 1 Introduction

The eukaryotic nuclear ribosomal ITS region (ITS1–5.8S–ITS2) provides high taxonomic resolution while being broadly amplifiable across fungi, supporting its adoption as a universal fungal barcode [Schoch et al., 2012]. In practice, amplicon reads often include conserved ribosomal flanks (SSU/18S and LSU/28S) alongside the variable ITS sequence. Retaining these conserved segments can bias similarity scoring and complicate taxonomic assignment, making extraction or trimming of ITS subregions a common preprocessing step [Bengtsson-Palme et al., 2013, Einarsson and Rivers, 2024].

ITSx identifies ITS boundaries using profile-HMM searches against conserved ribosomal domains, running HMMER with strict settings to maximize detection sensitivity [Bengtsson-Palme et al., 2013]. ITSxpress adapts this logic for FASTQ workflows, preserving quality scores and accelerating processing by reducing redundant HMM searches via dereplication or clustering [Rivers et al., 2018, Einarsson and Rivers, 2024].

Long-read amplicon sequencing generates large volumes of reads and presents practical challenges: throughput at scale, higher fractions of partial or truncated reads, and platform-specific error patterns that can affect boundary calling. An extractor designed for long reads can be valuable if it improves runtime and interpretability while maintaining boundary accuracy and downstream taxonomic performance comparable to established tools.

Here we describe ITSxRust, an ITS region extractor implemented in Rust, and evaluate it against ITSx and ITSxpress v2 on a real Oxford Nanopore ITS amplicon dataset.

## 2 Methods

### 2.1 Overview and design goals

ITSxRust is an ITS extractor implemented in Rust. The design preserves the established biological logic of HMM-based delimitation while targeting improved performance and pipeline integration for long-read datasets. The primary goals are to reduce overhead from process spawning and repeated text parsing, to enable streaming I/O with predictable memory use, and to provide richer per-read diagnostics than existing tools.

### 2.2 Architecture and implementation

ITSxRust takes a four-anchor chain approach to boundary detection. For each read, profile-HMM hits are classified into four anchor types, SSU 3^′^end, 5.8S 5^′^start, 5.8S 3^′^end, and LSU 5^′^start, on both strands. The tool retains the top-K hits per anchor type per strand (default *K* = 8) and enumerates all valid four-anchor chains (one hit per anchor type, all on the same strand) subject to configurable length constraints on the ITS1, ITS2, and full ITS intervals. The chain with the highest sum-of-scores is selected.

For reads where no valid four-anchor chain can be constructed, ITSxRust applies a partial-chain fallback: it attempts to delimit individual subregions using two-anchor pairs, SSU 3^′^end + 5.8S 5^′^start for ITS1, 5.8S 3^′^end + LSU 5^′^start for ITS2, and SSU 3^′^end + LSU 5^′^start for the full ITS region, subject to the same length constraints. This recovers reads that are missing one or two anchors (e.g., partial reads lacking the SSU flank) while preserving boundary quality, since each extracted region is still delimited by two profile-HMM hits.

Each extracted read is assigned a confidence label: **confident** (full four-anchor chain with all scores and E-values meeting thresholds), **partial** (from a two-anchor pair fallback), or **ambiguous** (chain found but at least one anchor below the score or E-value threshold). Reads for which no valid chain or pair can be constructed are reported with structured failure codes identifying the specific point of failure (e.g., missing anchor, constraint violation, coordinate conflict).

### 2.3 Input and output

ITSxRust reads single-end FASTQ or FASTA files (optionally gzip-compressed). Outputs include:

- Extracted sequences for ITS1, ITS2, and full ITS in FASTA or FASTQ format, with quality scores preserved when the input is FASTQ.
- A per-read TSV and/or JSONL report with anchor coordinates, scores, E-values, computed bounds, confidence labels, and failure reason codes.
- A per-sample JSON QC summary aggregating extraction counts (by confidence level), skip-reason breakdowns, and effective parameters, suitable for ingestion by pipeline aggregation tools such as MultiQC.
- Structured diagnostic messages for reads that fail extraction, identifying the stage of failure (no HMM hits, missing anchors, constraint violation, or coordinate conflict).

### 2.4 HMM search

ITSxRust performs profile-HMM searches using HMMER’s `nhmmer` program [Eddy, 2011]. The current implementation invokes `nhmmer` as an external process and parses the resulting `–tblout` output. To minimize overhead, ITSxRust streams the tblout file line-by-line, maintaining only the top-K hits per anchor type per strand per read in memory. Users can alternatively supply a pre-computed tblout file (e.g., from a previous run or a shared HMM search step in a pipeline), skipping the `nhmmer` invocation entirely.

### 2.5 Strand handling

Reads may be sequenced on either strand. ITSxRust normalizes all hit coordinates to a canonical forward orientation before chain selection. When the best chain is on the minus strand, the extracted subsequence is reverse-complemented (and quality scores reversed for FASTQ output) so that all output sequences are in the biological 5^′^ →3^′^ orientation.

### 2.6 Redundancy reduction

To reduce repeated HMM work on identical sequences, ITSxRust optionally performs exact dereplication: sequences are hashed and only unique representatives are searched, with results projected back to duplicates. This is conceptually aligned with the acceleration strategies of ITSxpress [Rivers et al., 2018, Einarsson and Rivers, 2024], adapted for the long-read setting.

### 2.7 Platform presets

ITSxRust provides parameter presets for common long-read platforms:

- `–preset ont`: relaxed length constraints and E-value thresholds to accommodate higher error rates and partial reads typical of Oxford Nanopore data.
- –`preset hifi`: stricter thresholds reflecting the higher accuracy of PacBio HiFi reads.

Users can override individual parameters when using a preset.

### 2.8 Benchmark dataset

We benchmarked all three tools on a publicly available Oxford Nanopore ITS amplicon dataset (SRA accession SRR21494940; 54,659 reads). All tools used the ITSx fungal HMM profile set (F.hmm, ITSx version 1.1.3) and were run with 8 threads on the same hardware (Apple M-series, macOS). ITSxRust was tested in two configurations: default parameters (*E* = 10^−5^, length constraints ITS1 50–1500 bp, ITS2 50–2000 bp, full 150–4000 bp) and ONT preset (*E* = 10^−3^, relaxed constraints ITS1 30–1800 bp, ITS2 30–2500 bp, full 100–5000 bp). ITSx was run with `-t F –save_regions all –detailed_results T`. ITSxpress v2 was run separately for ITS1, ITS2, and ALL regions with `–taxa Fungi –single_end`.

For boundary accuracy and classification evaluation, we selected 2,000 reads for which ITSx identified all five ribosomal regions (SSU, ITS1, 5.8S, ITS2, LSU) as ground-truth references with known ITS coordinates. We simulated 14,751 ONT-like reads from these references using Badread [Wick, 2019] (identity 92–98%, mean read length 800 bp). Ground-truth taxonomy was established by classifying the error-free reference sequences against the UNITE database (dynamic release 2025-02-19) [Nilsson et al., 2024] using VSEARCH [Rognes et al., 2016] at 80% identity.

### 2.9 Comparators and evaluation metrics

Comparators are ITSx [Bengtsson-Palme et al., 2013] and ITSxpress v2 [Rivers et al., 2018, Einarsson and Rivers, 2024]. Metrics:

- **Throughput:** wall time and peak resident set size (RSS), measured using GNU time.
- **Extraction success:** proportion of reads yielding ITS1, ITS2, and full ITS extracts, including the contribution of partial-chain fallback in ITSxRust.
- **Boundary accuracy (simulation):** each tool’s extracted sequences are aligned back to reference sequences with minimap2; alignment boundaries on the reference are compared to the known true ITS coordinates.
- **Downstream classification:** genus- and species-level agreement between each tool’s classified extractions and the classification of the error-free reference sequences.

**Figure 1:**
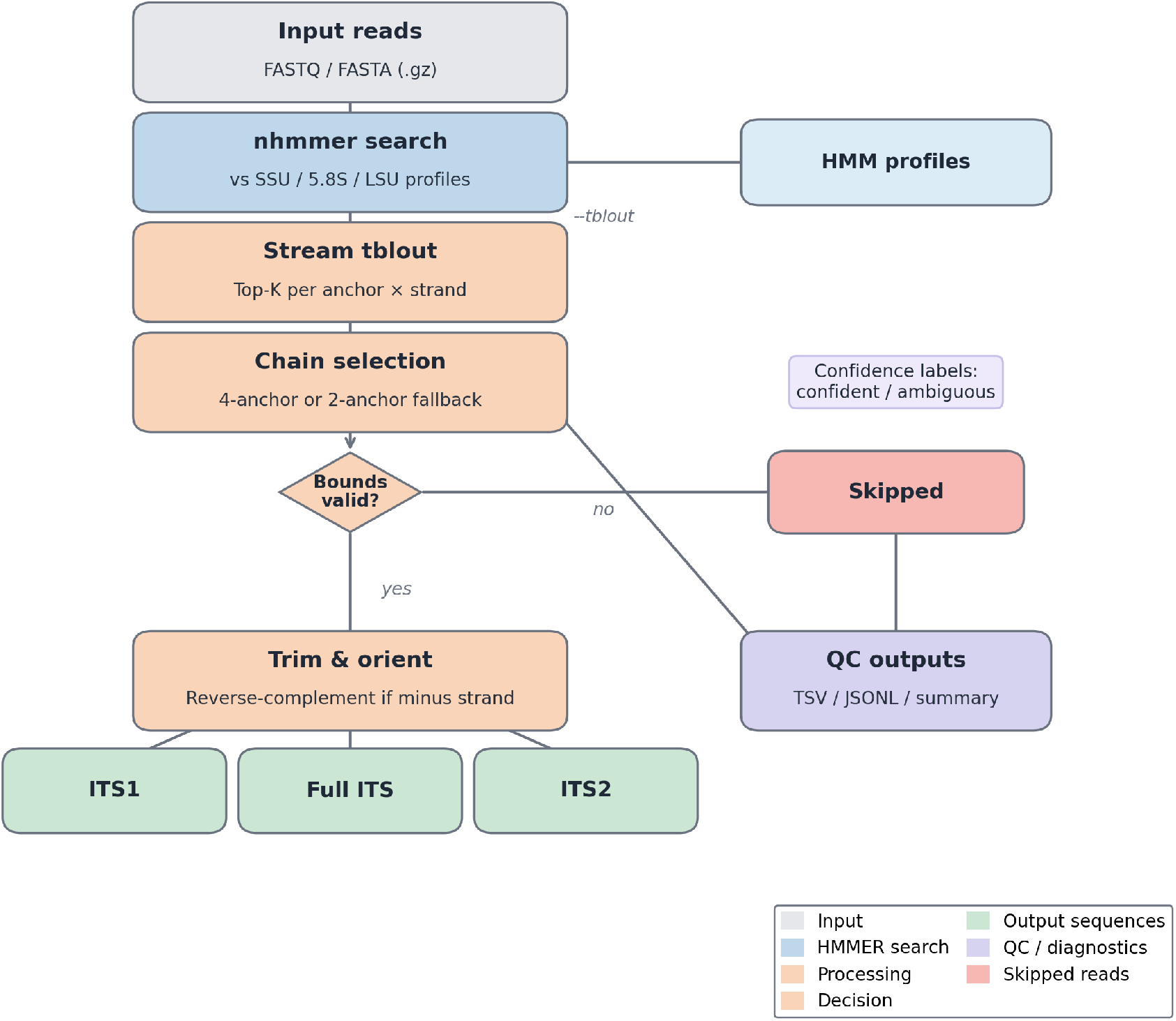
ITSxRust pipeline overview. Input reads are searched against SSU, 5.8S, and LSU profile HMMs using `nhmmer`. The tblout is streamed to retain the top-K hits per anchor per strand per read. Chains are formed (4-anchor or 2-anchor fallback) and classified as confident or ambiguous. Valid chains are trimmed to the requested ITS region(s) and reverse-complemented if on the minus strand. Reads that cannot form a valid chain are reported with structured skip reasons.

## 3 Results

### 3.1 Feature comparison

Table 1 summarizes key design differences among the three tools. ITSxRust is designed for longread workflows, with streaming tblout parsing, a four-anchor chain model with two-anchor partial fallback, per-read confidence labels, and platform-specific parameter presets.

**Table 1:** Feature comparison among ITS extractors.

| Capability | ITSx | ITSxpress v2 | ITSxRust |
| --- | --- | --- | --- |
| Primary input | FASTA | FASTQ | FASTQ and FASTA |
| Quality preservation | No | Yes | Yes |
| HMMER usage | External | External | External + streaming parse |
| Redundancy reduction | None | Cluster/derep | Exact dereplication |
| Boundary strategy | Flexible (partial OK) | Full profile | 4-anchor chain + 2-anchor fallback |
| Long-read emphasis | General | Noted | ONT/HiFi presets |
| Confidence labels | No | No | Confident / Partial / Ambiguous |
| Failure reporting | Limited | Limited | Per-read codes + JSON QC summary |

### 3.2 Throughput and resource use

Table 2 reports wall time and peak memory for each tool on the ONT dataset (54,659 reads, 8 threads). ITSxRust completed in approximately 15 minutes, 4.6 × faster than ITSx (72 minutes). ITSxpress v2 was the fastest tool at 7–8 minutes, reflecting its clustering-based redundancy reduction; however, this speed came at a substantial cost in extraction yield (Section 3.3).

**Table 2:**
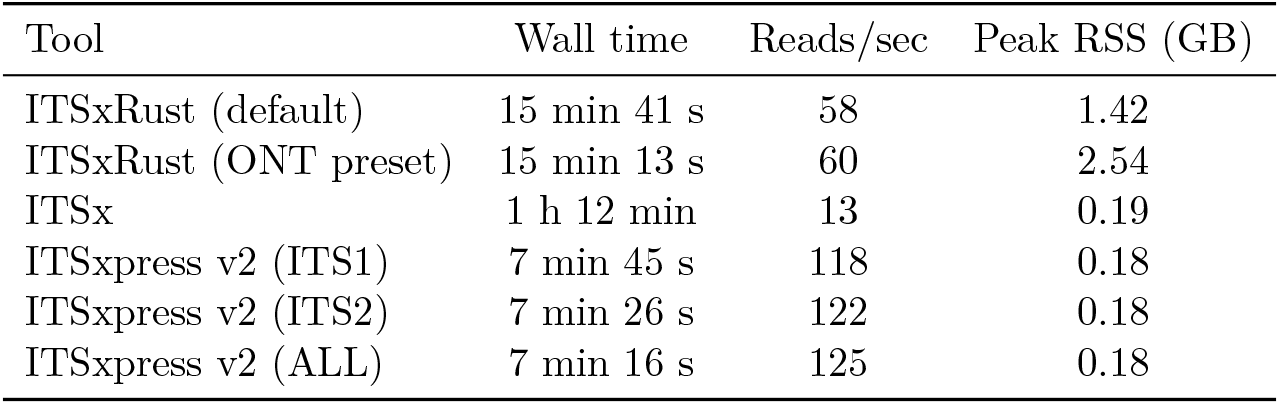
Throughput and resource use on the ONT dataset (54,659 reads, 8 threads).

| Tool | Wall time | Reads/sec | Peak RSS (GB) |
| --- | --- | --- | --- |
| ITSxRust (default) | 15 min 41 s | 58 | 1.42 |
| ITSxRust (ONT preset) | 15 min 13 s | 60 | 2.54 |
| ITSx | 1 h 12 min | 13 | 0.19 |
| ITSxpress v2 (ITS1) | 7 min 45 s | 118 | 0.18 |
| ITSxpress v2 (ITS2) | 7 min 26 s | 122 | 0.18 |
| ITSxpress v2 (ALL) | 7 min 16 s | 125 | 0.18 |

ITSxRust’s higher memory footprint (1.4–2.6 GB vs. ∼ 0.2 GB) reflects `nhmmer` loading the full sequence set into memory for the nucleotide search.

### 3.3 Extraction success

Table 3 shows the proportion of reads from which each tool successfully extracted ITS1, ITS2, and full ITS sequences. ITSxRust with the ONT preset achieved 93.1% ITS2 extraction and 75.3% full ITS extraction, the latter exceeding ITSx’s 69.9%. The partial-chain fallback contributed 10,725 additional reads (19.6% of the dataset) that lacked a complete four-anchor chain but had sufficient anchors for at least one subregion.

**Table 3:**
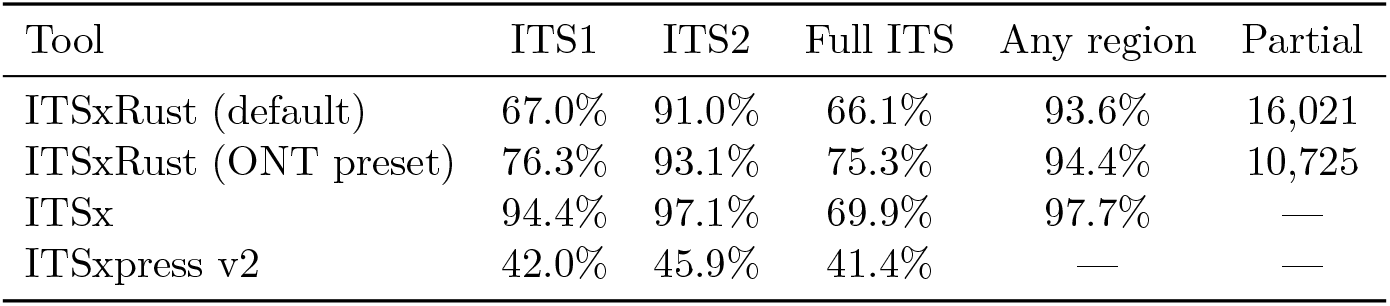
Extraction success rates on the ONT dataset (54,659 reads). Percentages are of total reads. ITSxRust rows include both four-anchor chain and partial-chain extractions; the partial contribution is shown separately.

ITSx achieved the highest per-region rates (94.4% ITS1, 97.1% ITS2) because it can delimit regions using flexible boundary strategies including read-edge fallback for truncated reads. ITSxpress v2 extracted substantially fewer reads (42–46%), reflecting its clustering-based approach: on ONT data where each read has unique error patterns, almost no sequences cluster, effectively disabling the dereplication benefit and causing many reads to be dropped.

The remaining gap in ITS1 extraction (76.3% vs. ITSx’s 94.4%) is primarily due to reads that begin within the ITS1 region, lacking the SSU 3^′^end anchor needed as a left delimiter. ITSx handles these by using the read boundary as a fallback left delimiter; ITSxRust currently requires a profile-HMM hit at both boundaries.

Table 4 shows the structured failure-reason breakdown from ITSxRust’s QC summary for reads that could not be extracted. The dominant failure mode is missing anchors, reads lacking one or more of the four anchor types, accounting for the majority of skipped reads. This diagnostic information is unique to ITSxRust and immediately actionable: a high rate of missing SSU 3^′^end anchors indicates that amplicon primers are cutting into the ITS1 region, which guides primer design and experimental troubleshooting.

**Table 4:** ITSxRust skip-reason breakdown (ONT preset, 54,659 reads).

| Skip reason | Reads |
| --- | --- |
| Missing anchors (partial read, no SSU/LSU hit) | 2,672 |
| No valid chain or pair under constraints | 374 |
| No HMM hits at all | 0 |
| Total skipped | 3,046 (5.6%) |

**Figure 2:**
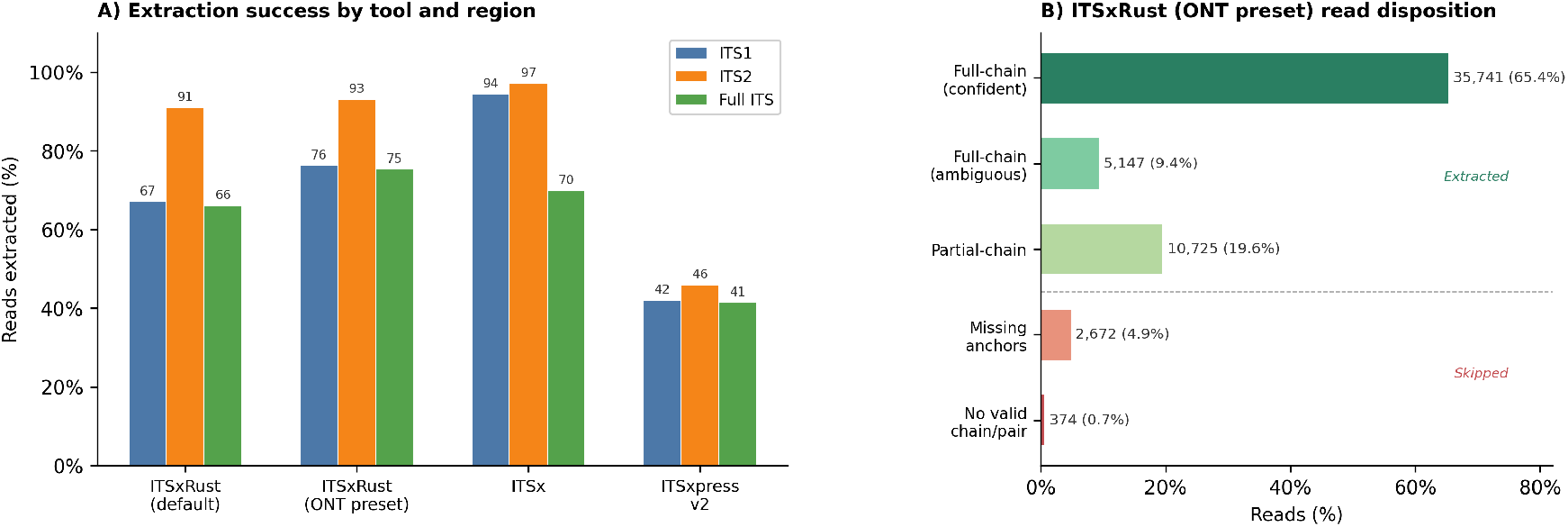
Extraction success on the ONT dataset (54,659 reads). **(A)** Grouped bar chart of extraction rates by tool and ITS region. **(B)** ITSxRust (ONT preset) read disposition, showing the contribution of full-chain (confident and ambiguous) and partial-chain extractions, and the breakdown of skipped reads by failure reason.

### 3.4 Boundary accuracy on simulated reads

To evaluate boundary accuracy, we used 2,000 reads from the ONT dataset for which ITSx identified all five ribosomal regions as ground-truth references with known ITS coordinates. We simulated 14,751 ONT-like reads from these references using Badread [Wick, 2019] (mean identity 92–98%, mean length 800 bp), ran all three tools on the simulated reads, then aligned each tool’s extracted sequences back to the original references with minimap2. The alignment boundaries on the reference define the tool’s called ITS coordinates, which are compared to the known truth.

Table 5 reports boundary accuracy per tool and region. For the full ITS region, all three tools achieved comparable accuracy: median absolute errors of 8–24 bp and 43–50% of reads within *±* 10 bp. For ITS1 and ITS2 evaluated separately, median errors are higher because many simulated reads are partial fragments that do not span the full reference; when the true boundary falls outside the read, the tool either does not extract the region or calls a boundary at the read edge. Among reads that were extracted, ITSxRust achieved ITS2 boundaries within ≤ 5 bp for 44.3% of reads, compared to 33.7% for ITSx and 47.2% for ITSxpress.

**Table 5:** Boundary accuracy on simulated ONT reads (14,751 reads from 2,000 references, Badread 92–98% identity). Maximum of start and end absolute errors is reported per read. *N* = number of reads extracted and successfully aligned to a reference.

| Tool | Region | Med. abs. err. (bp) | Mean | 95th %ile | % $\leq 5$ bp |
| --- | --- | --- | --- | --- | --- |
| ITSxRust | ITS1 | 70 | 211 | 425 | 36.9% |
| ITSxRust | ITS2 | 11 | 258 | 538 | 44.3% |
| ITSxRust | Full ITS | 14 | 18 | 33 | 43.4% |
| ITSx | ITS1 | 292 | 238 | 447 | 18.7% |
| ITSx | ITS2 | 119 | 269 | 559 | 33.7% |
| ITSx | Full ITS | 24 | 17 | 32 | 44.3% |
| ITSxpress v2 | ITS1 | 17 | 205 | 425 | 39.3% |
| ITSxpress v2 | ITS2 | 7 | 245 | 537 | 47.2% |
| ITSxpress v2 | Full ITS | 8 | 17 | 33 | 46.3% |

**Figure 3:**
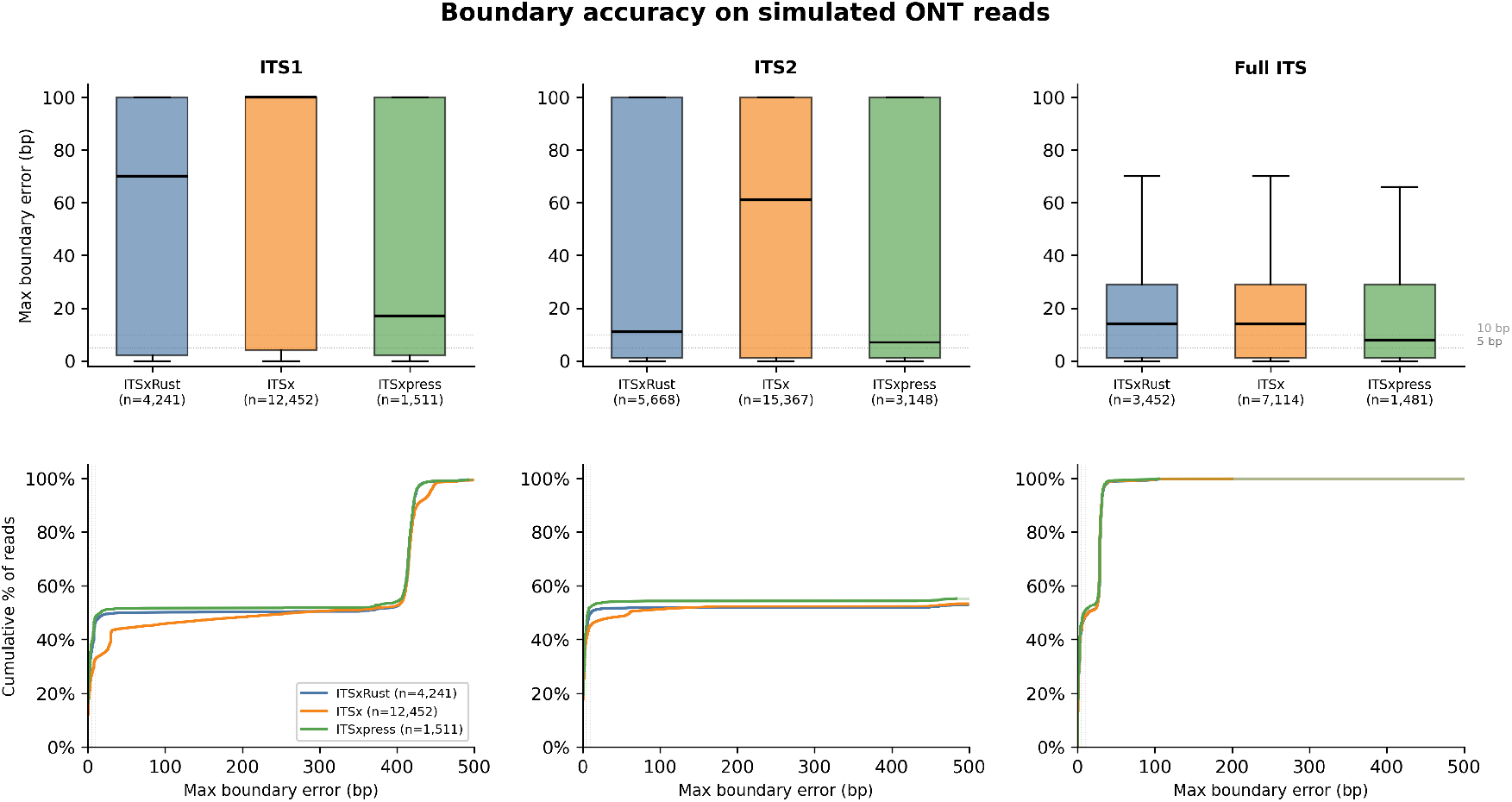
Boundary error distributions on simulated ONT reads (14,751 reads from 2,000 references). Top row: Box plots of the maximum of start/end absolute error per read, capped at 100 bp for readability (outliers hidden). Bottom row: Empirical cumulative distribution functions (ECDFs) up to 50 bp.

Table 6 reports the same evaluation on simulated HiFi reads (15,000 reads, 99–100% identity). As expected, higher read accuracy improved boundary precision across all tools: ITSxRust achieved a median full ITS error of 3 bp (51.0% within ≤ 5 bp), comparable to ITSx (2 bp, 51.3%) and ITSxpress (10 bp, 49.8%). The ITS1 and ITS2 subregion errors remained elevated for ITSx and ITSxpress due to the same partial-fragment effect observed in the ONT simulation, while ITSxRust showed notably lower ITS1 median error (9 bp vs. 265–318 bp) reflecting its stricter two-anchor requirement filtering out reads with edge-called boundaries.

**Table 6:** Boundary accuracy on simulated HiFi reads (15,000 reads subsampled from 165,215; 2,000 references). Maximum of start and end absolute errors is reported per read. *N* = number of reads extracted and successfully aligned to a reference.

| Tool | Region | Med. abs. err. (bp) | Mean | 95th %ile | % $\leq 5$ bp |
| --- | --- | --- | --- | --- | --- |
| ITSxRust | ITS1 | 9 | 203 | 421 | 48.4% |
| ITSxRust | ITS2 | 3 | 257 | 535 | 51.0% |
| ITSxRust | Full ITS | 3 | 17 | 31 | 51.0% |
| ITSx | ITS1 | 318 | 237 | 424 | 31.8% |
| ITSx | ITS2 | 135 | 266 | 538 | 40.3% |
| ITSx | Full ITS | 2 | 17 | 31 | 51.3% |
| ITSxpress v2 | ITS1 | 265 | 211 | 421 | 48.5% |
| ITSxpress v2 | ITS2 | 3 | 258 | 535 | 51.0% |
| ITSxpress v2 | Full ITS | 10 | 16 | 31 | 49.8% |

**Figure 4:**
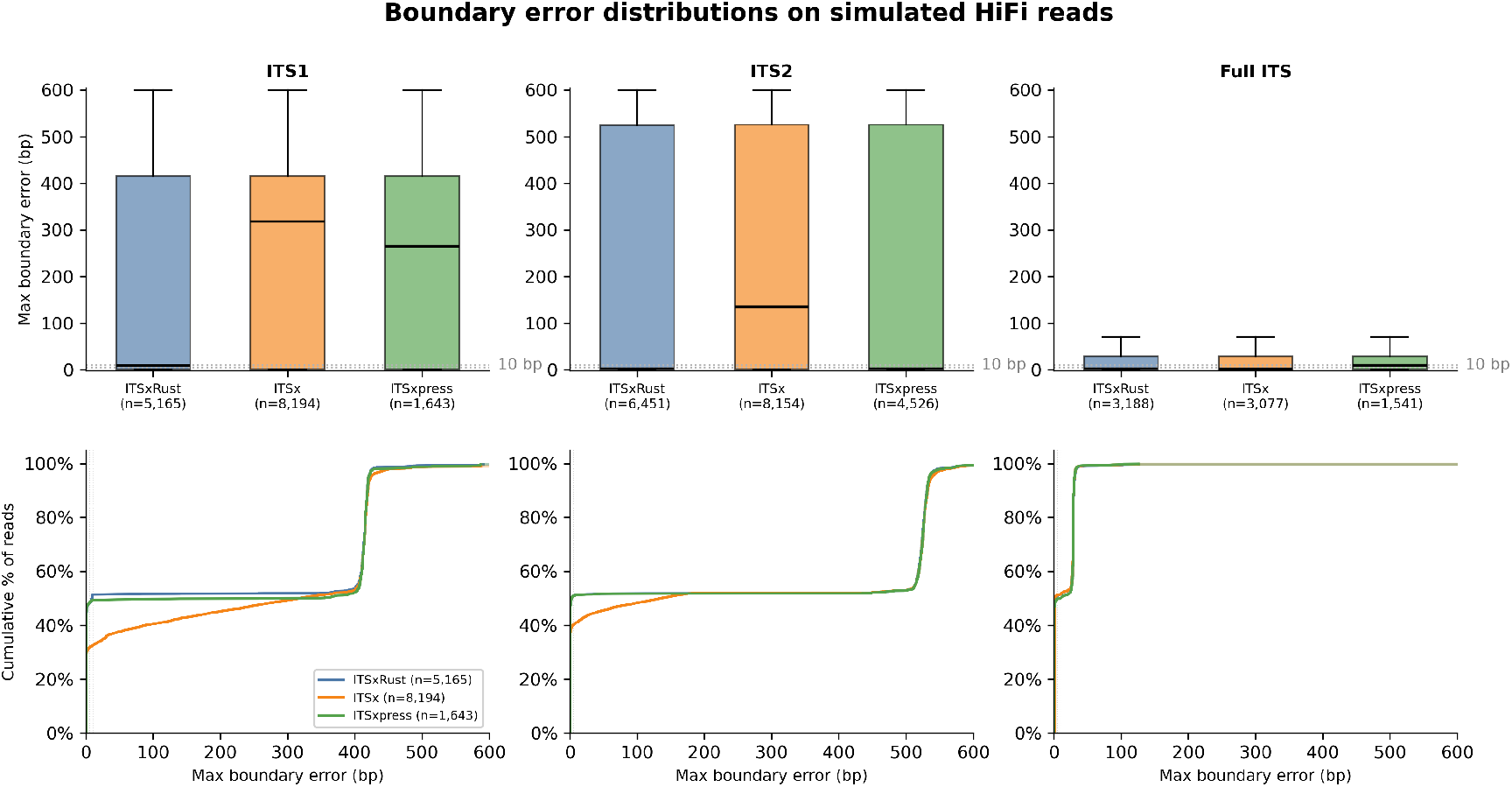
Boundary error distributions on simulated HiFi reads. Boundary accuracy was evaluated on 15,000 simulated HiFi reads subsampled from a larger dataset. For each extracted read, the maximum of the absolute start and end boundary errors is shown. Top row: boxplots of perread maximum boundary error (clipped at 600 bp for visualization). Bottom row: cumulative percentage of reads as a function of maximum boundary error. Sample sizes (*n*) indicate the number of extracted reads successfully aligned to a reference.

### 3.5 Downstream classification impact

To assess whether ITS extraction affects taxonomic assignment, we classified the simulated reads against the UNITE database (dynamic release 2025-02-19) [Nilsson et al., 2024] using VSEARCH [Rognes et al., 2016] at 80% identity. Ground-truth taxonomy was established by classifying the error-free reference sequences; we then measured genus- and species-level agreement between each tool’s extracted reads and the corresponding reference classification.

Table 7 shows that all three extraction tools substantially improved classification accuracy relative to raw (untrimmed) reads, confirming that removing conserved flanks reduces spurious similarity matches. Genus accuracy increased from 95.2% (raw) to 98.3–99.8% after trimming, and species accuracy from 94.1% to 98.3–99.7%. ITSxRust achieved the highest per-region classification accuracy for ITS1 (99.7% genus) and matched ITSx and ITSxpress on ITS2 and full ITS. The modest differences among tools (*<*2% across all comparisons) indicate that all three produce biologically equivalent extracted sequences for downstream classification.

**Table 7:** Classification accuracy on simulated ONT reads using UNITE and VSEARCH. Accuracy is measured as agreement with the classification of the error-free reference sequence. *N* = reads classified against UNITE that could be traced to a reference.

| Input to classifier | $N$ | Genus acc. | Species acc. |
| --- | --- | --- | --- |
| Raw (untrimmed) | 3,249 | 95.2% | 94.1% |
| ITSxRust (full ITS) | 1,666 | 99.7% | 99.6% |
| ITSxRust (ITS1) | 2,044 | 99.7% | 99.5% |
| ITSxRust (ITS2) | 2,912 | 99.2% | 99.2% |
| ITSx (full ITS) | 1,033 | 99.8% | 99.7% |
| ITSx (ITS1) | 3,389 | 98.5% | 98.3% |
| ITSx (ITS2) | 3,528 | 98.3% | 98.3% |
| ITSxpress v2 (full ITS) | 744 | 99.7% | 99.6% |
| ITSxpress v2 (ITS1) | 746 | 99.5% | 99.3% |
| ITSxpress v2 (ITS2) | 1,629 | 99.2% | 99.2% |

HiFi simulation results (Table 8) showed a similar pattern: all tools achieved ≥ 98.1% genus accuracy after trimming, with inter-tool differences below 2%. Raw untrimmed HiFi reads classified at 94.8% genus accuracy, confirming that flank removal benefits classification regardless of read accuracy.

**Table 8:** Classification accuracy on simulated HiFi reads using UNITE and VSEARCH. Accuracy is measured as agreement with the classification of the error-free reference sequence. *N* = reads classified against UNITE that could be traced to a reference.

| Input to classifier | $N$ | Genus acc. | Species acc. |
| --- | --- | --- | --- |
| Raw (untrimmed) | 3,217 | 94.8% | 94.7% |
| ITSxRust (full ITS) | 1,651 | 99.8% | 99.8% |
| ITSxRust (ITS1) | 2,685 | 99.5% | 99.4% |
| ITSxRust (ITS2) | 3,338 | 98.9% | 98.9% |
| ITSx (full ITS) | 1,595 | 99.7% | 99.7% |
| ITSx (ITS1) | 4,296 | 98.4% | 98.3% |
| ITSx (ITS2) | 4,281 | 98.1% | 98.1% |
| ITSxpress v2 (full ITS) | 775 | 99.9% | 99.9% |
| ITSxpress v2 (ITS1) | 834 | 98.6% | 98.3% |
| ITSxpress v2 (ITS2) | 2,323 | 99.0% | 99.0% |

## 4 Discussion

On an Oxford Nanopore ITS dataset of 54,659 reads, ITSxRust achieved 4.6 × the throughput of ITSx while extracting a comparable or higher proportion of reads across ITS subregions. The full ITS extraction rate (75.3% with the ONT preset) exceeded ITSx’s 69.9%, and ITS2 extraction (93.1%) approached ITSx’s 97.1%. Both tools substantially outperformed ITSxpress v2, which extracted only 42–46% of reads on this long-read dataset.

The partial-chain fallback proved critical for competitive extraction rates. Without it, ITSxRust’s four-anchor chain requirement yielded only 64–75% of reads, since many ONT amplicon reads are truncated and lack one or more conserved flanks. The two-anchor pair fallback recovered an additional 10,725 reads (19.6%), bringing the overall extraction rate to 94.4% (any region) while maintaining boundary quality: every extracted region is still delimited by two profile-HMM hits rather than arbitrary read boundaries.

The structured diagnostic output is a practical contribution for long-read workflows. The QC summary revealed that the dominant failure mode was missing anchors (specifically SSU 3^′^end), immediately indicating that many reads in this dataset begin within the ITS1 region rather than upstream in the SSU. This kind of diagnostic, unavailable from ITSx or ITSxpress, allows users to distinguish between biological causes of extraction failure (e.g., primer placement) and technical ones (e.g., HMMER sensitivity, constraint settings), enabling targeted troubleshooting at the batch level.

ITSxpress v2’s low extraction rate on this dataset reflects a fundamental mismatch between its clustering-based acceleration strategy and long-read error profiles. Each ONT read carries a unique pattern of errors, so almost no sequences cluster; the dereplication step provides negligible speedup while the overall pipeline drops the majority of reads. ITSxpress was designed for short-read Illumina amplicons where identical or near-identical sequences are common, and its performance on that platform is well-established [Rivers et al., 2018, Einarsson and Rivers, 2024].

ITSxRust’s exact dereplication option (`–derep`) faces a similar limitation on ONT data, where per-read error patterns make identical sequences rare; it is expected to provide greater benefit on PacBio HiFi amplicon datasets, where higher read accuracy produces more exact-duplicate sequences.

The remaining ITS1 extraction gap between ITSxRust (76.3%) and ITSx (94.4%) is a known design trade-off. ITSx can delimit ITS1 using the read boundary as a left edge when the SSU flank is absent, whereas ITSxRust requires a profile-HMM hit on both sides of the extracted region. Supporting read-edge fallback is feasible and would close this gap, but introduces reads with less certain left boundaries; we leave this as a user-configurable option for a future release.

ITSxRust takes a complementary approach to acceleration compared to prior tools: rather than reducing the number of HMMER searches via clustering, it focuses on reducing the overhead surrounding the search, streaming tblout parsing, parallel chain computation, and Rust-native I/O. The current bottleneck is `nhmmer` itself (*>*99% of wall time), so future in-process integration via FFI bindings to libhmmer or exploration of alternative HMM search engines could provide further speedups.

On simulated ONT reads with known true ITS boundaries, all three tools achieved comparable boundary accuracy for the full ITS region (median errors of 8–24 bp; Table 5). Per-region errors (ITS1, ITS2) were higher across all tools because many simulated reads are partial fragments that do not span the full reference, inflating error statistics when the true boundary falls outside the read. ITSx showed notably higher ITS1 median error (292 bp vs. 17–70 bp for ITSxpress and ITSxRust) because its read-edge fallback strategy places the left ITS1 boundary at position 1 for truncated reads; while this maximizes extraction yield (94.4% on real data), it shifts boundary coordinates substantially when the read begins mid-ITS1.

Classification accuracy against UNITE was uniformly high after extraction by any tool (≥ 98.3% genus accuracy; Table 7), and all tools improved upon raw untrimmed reads (95.2%). The small inter-tool differences (*<*2%) suggest that boundary accuracy variations at the level observed here do not materially affect downstream taxonomic assignment, and the choice of extractor can be driven by practical considerations such as speed, diagnostic output, and extraction yield rather than classification fidelity.

### 4.1 Limitations and future work

Several limitations should be noted:

- **External HMMER dependency:** The current implementation invokes `nhmmer` as a subprocess. In-process integration via Rust FFI bindings to libhmmer would eliminate process startup overhead and intermediate I/O, and is planned for a future release.
- **ITS1 extraction on truncated reads:** ITSxRust requires at least SSU 3^′^end and 5.8S 5^′^start anchors to delimit ITS1. Reads that begin within the ITS1 region (lacking the SSU flank) cannot be extracted, whereas ITSx can use the read boundary as a left delimiter. Supporting read-edge fallback boundaries is a potential future enhancement.
- **Profile HMM dependence:** Results depend on the quality and taxonomic breadth of the HMM profiles used. ITSxRust is tested with the ITSx fungal profiles (F.hmm) but should be validated with other organism groups.
- **Clustering-induced boundary drift:** The optional dereplication step groups only exact duplicates; approximate clustering for error-tolerant grouping (as in ITSxpress) is not yet implemented and, if added, would require careful validation against simulation data.

## 5 Availability and reproducibility

- **Source code:** https://github.com/ayobi/itsxrust
- **Releases:** Semantic versioning with tagged benchmark scripts.
- **Installation:** Bioconda recipe and prebuilt binaries; container images (Docker).
- **Reproducibility:** Benchmark datasets (or accessions), exact commands, and environment specifications are provided in the repository.

## Acknowledgements

Public sequencing data were obtained from NCBI SRA (accession SRR21494940). I also acknowledge Bengtsson-Palme et al. for the ITSx HMM profiles used by ITSxRust, and the Bioconda community for packaging support.

## Author contributions

A.O. designed and developed the software, conducted benchmarking experiments, analyzed results, and wrote the manuscript. C.L., K.F., and B.O. contributed to discussions on full-length ITS sequencing as part of ongoing research at the Centro de Biotecnología de Sistemas, with C.L. coordinating sequencing efforts and K.F. and B.O. conducting library preparation and sequencing. P.P. supervised the research and provided critical feedback on the manuscript.

